# *N*^6^-methyladenosine (m^6^A) and reader protein YTHDF2 enhance innate immune response by mediating DUSP1 mRNA degradation and activating mitogen-activated protein kinases during bacterial and viral infections

**DOI:** 10.1101/2022.12.01.518805

**Authors:** Jian Feng, Wen Meng, Luping Chen, Xinquan Zhang, Ashley Markazi, Weiming Yuan, Yufei Huang, Shou-Jiang Gao

**Author notes:** Address correspondence to Shou-Jiang Gao,.

## Abstract

Mitogen-activated protein kinases (MAPKs) play critical roles in the induction of numerous cytokines, chemokines, and inflammatory mediators that mobilize the immune system to counter pathogenic infections. Dual-specificity phosphatase-1 (DUSP1) is a member of dual-specificity phosphatases, which inactivates MAPKs through a negative feedback mechanism. Here we report that in response to viral and bacterial infections, not only DUSP1 transcript but also its *N*^6^-methyladenosine (m^6^A) level rapidly increase together with the m^6^A reader protein YTHDF2, resulting in enhanced YTHDF2-mediated DUSP1 transcript degradation. Knockdown of DUSP1 promotes p38 and JNK phosphorylation and activation, thus increasing the expression of innate immune response genes including IL1β, CSF3, TGM2 and SRC. Similarly, knockdown of m^6^A eraser ALKBH5 increases DUSP1 transcript m^6^A level resulting in accelerated transcript degradation, activation of p38 and JNK, and enhanced expression of IL1β, CSF3, TGM2 and SRC. These results demonstrate that m^6^A and reader protein YTHDF2 orchestrate optimal innate immune response during viral and bacterial infections by downregulating the expression of a negative regulator DUSP1 of the p38 and JNK pathways that are central to innate immune response against pathogenic infections.

**IMPORTANCE:** Innate immunity is central for controlling pathogenic infections and maintaining the homeostasis of the host. In this study, we have revealed a novel mechanism regulating innate immune response during viral and bacterial infections. We have found that *N*^6^-methyladenosine (m^6^A) and the reader protein YTHDF2 regulate dual-specificity phosphatase-1, a negative regulator of mitogen-activated protein kinases p38 and JNK, to maximize innate immune response during viral and bacterial infections. These results provide novel insights into the mechanism regulating innate immunity, which could help the development of novel approaches for controlling pathogenic infections.

## Introduction

The innate immune system is a highly efficient cellular and molecular network in mammalian cells that protects the organism against pathogenic infections (1). This first line of defense against invasion is achieved by sensing the pathogens through pattern recognition receptors (2). Stimulation of pattern recognition receptors on the cell surface and in the cytoplasm of innate immune cells activates multiple mitogen-activated protein kinases (MAPKs) including the extracellular signal-regulated kinase (ERK), p38 and Jun N-terminal kinase (JNK) (3). MAPKs are a group of highly conserved serine/threonine protein kinases in eukaryotes (4), which play a critical role in inducing numerous cytokines, chemokines, and inflammatory mediators that mobilize the immune system to counter pathogenic infections (5). Furthermore, the induction of pro-inflammatory response promotes the recruitment of additional immune cells to invoke secondary innate and adaptive immune responses (6).

Dual-specificity phosphatase-1 (DUSP1, also known as MAPK phosphatase-1 or MKP-1) was initially identified in cultured murine cells (7). It is a member of dual-specificity phosphatases (DUSPs), which are key players for inactivating different MAPKs (8). DUSP1 expression is enhanced upon numerous pathogenic infections, and it is an important feedback mechanism for controlling excessive immune response and inflammation (9, 10). By dephosphorylation, DUSP1 inhibits the activation of specific threonine and tyrosine residues on p38 and JNK, resulting in inactivation of inflammatory or innate immune response through inhibiting the expression of numerous effector genes at transcriptional or post-transcriptional levels (11).

*N6*-methyladenosine (m^6^A), a dynamic posttranscriptional RNA modification, is critical for almost all aspects of RNA metabolism and functions including structure, maturation, stability, splicing, export, translation, and decay (12). Recent studies show that m^6^A modification not only directly regulates the expression of innate immune response genes, but also indirectly affects the mRNA metabolism pathway to further regulate the innate immune response during bacterial and viral infections (13–16).

We have previously shown that m^6^A plays an important role in regulating innate immune response against both bacterial and viral infections by directly and indirectly regulating the expression of innate immune response genes (13). More recent works indicate m^6^A is a vital factor for regulating innate immune response and cytokines by affecting the IKKε/TBK1/IRF3, MAPK and NF-κB pathways (17, 18). In this study, we have discovered that DUSP1 is a direct m^6^A target, and m^6^A and the reader protein YTHDF2 regulate DUSP1 stability to maximize innate immune response during bacterial and viral infections.

## Results

### m^6^A mediates DUSP1 transcript expression during bacterial infection

We have previously mapped the cellular expression profiles and m^6^A epitranscriptomes, and identified a set of genes including innate immune response genes that are differentially methylated and differentially expressed during viral and bacterial infections (13). Among them, DUSP1, an important regulator of innate immune response genes, was significantly hyper-methylated during gram-negative bacteria *Pseudomonas aeruginosa* infection, which peaked at 2 h post-infection (hpi), then decreased at 4 and 6 hpi (Fig. 1A). At the same time, DUSP1 transcript expression was upregulated which also peaked at 2 hpi, then decreased at 4 and 6 hpi (Fig. 1B). These results were consistent with the induction of DUSP1 by LPS or TLR ligands reported in previous studies (19, 20).

**FIG 1.**
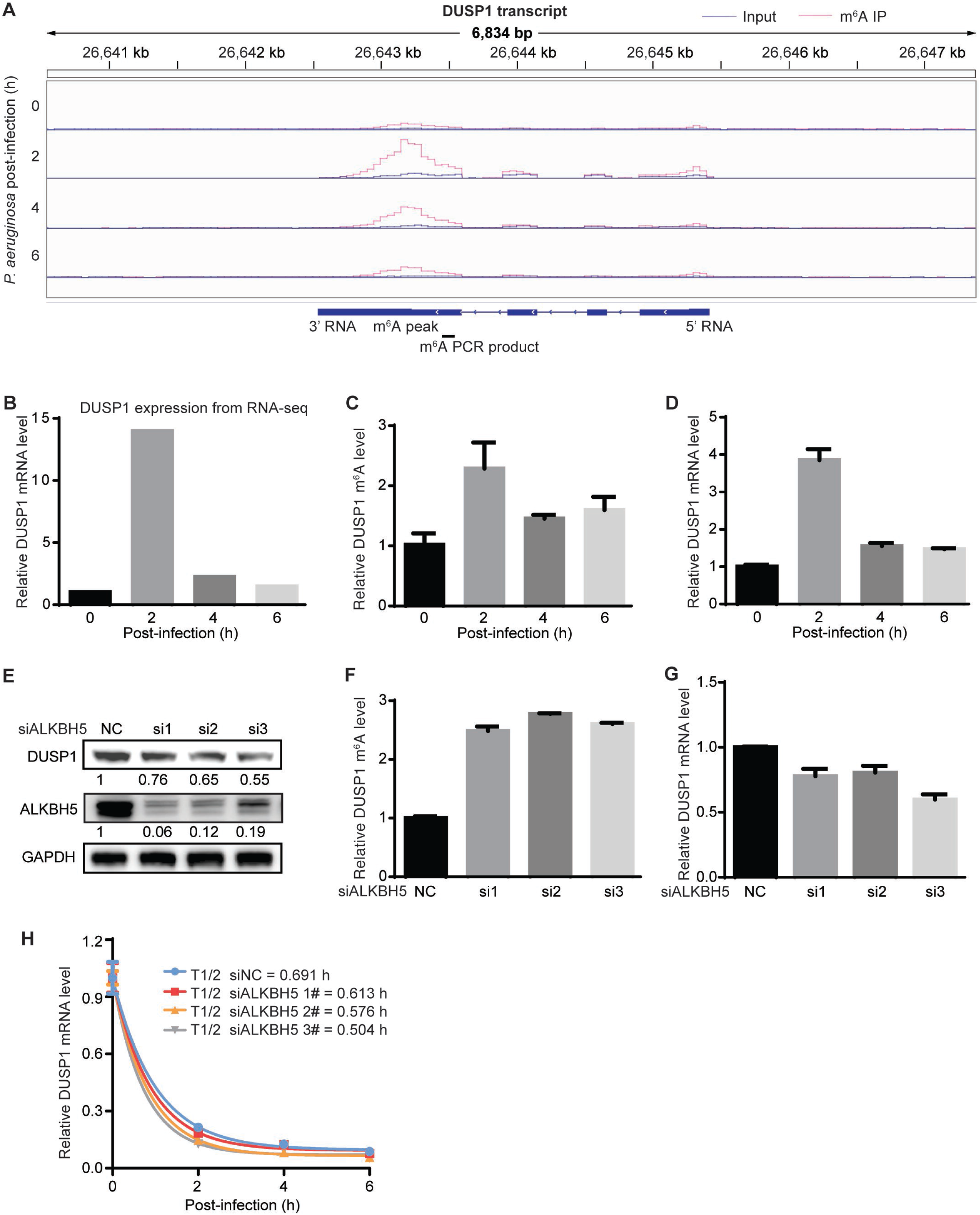
m^6^A mediates DUSP1 transcript stability during bacterial infection. (A) Tracks of m^6^A peaks on DUSP1 transcript at 0, 2, 4, and 6 hpi of *P. aeruginosa*.(B) Expression levels of DUSP1 transcript at 0, 2, 4, and 6 hpi of *P. aeruginosa* quantified by RNA-seq. (C) m^6^A levels on DUSP1 transcript at 0, 2, 4, and 6 hpi of *P. aeruginosa* examined by MeRIP-qPCR. (D) Expression levels of DUSP1 transcript at 0, 2, 4, and 6 hpi of *P. aeruginosa* quantified by RT-qPCR. (E) Examination of DUSP1 and ALKBH5 protein levels following ALKBH5 knockdown in RAW264.7 cells by Western-blotting. (F) m^6^A levels on DUSP1 transcript following ALKBH5 knockdown in RAW264.7 cells examined by MeRIP-qPCR. (G) Expression levels of DUSP1 transcript following ALKBH5 knockdown in RAW264.7 cells examined by RT-qPCR. (H) Alterations of half-lives of DUSP1 transcript following ALKBH5 knockdown in RAW264.7 cells during *P. aeruginosa* infection examined by RT-qPCR at the indicated time points following addition of 10 μg/ml actinomycin D.

We confirmed the increase of DUSP1 transcript m^6^A during *P. aeruginosa* infection by m^6^A-immunoprecipitation reverse transcription quantitative real time PCR (MeRIP-qPCR). The DUSP1 transcript m^6^A level was increased by 2.3-fold at 2 hpi of *P. aeruginosa* but then decreased at 4 and 6 hpi (Fig. 1C). Reverse transcription quantitative real time PCR (RT-qPCR) further confirmed the increased DUSP1 transcript expression following *P. aeruginosa* infection, which peaked at 2 hpi (Fig. 1D).

We then performed knockdown of ALKBH5, an m^6^A “eraser”, to determine whether the increase of DUSP1 transcript m^6^A could affect its expression (Fig. 1E). As expected, ALKBH5 knockdown further increased the m^6^A level of DUSP1 transcript by 2.5- to 2.8-fold (Fig. 1F). However, DUSP1 transcript expression was reduced by 25% to 40% (Fig. 1G), which was also reflected in the decreased DUSP1 protein level (Fig. 1E). These results suggest that the increased m^6^A level during *P. aeruginosa* infection likely serves to reverse the upregulation of DUSP1 transcript. Since one of the functions of m^6^A modification is to mediate RNA decay (21, 22), we examined DUSP1 transcript stability during *P. aeruginosa* infection. The half-life of DUSP1 transcript was reduced by 12.7% to 37.1% following ALKBH5 knockdown (Fig. 1H), indicating m^6^A regulation of DUSP1 RNA decay during *P. aeruginosa* infection.

### YTHDF2 mediates m^6^A-dependent DUSP1 transcript degradation

In order to further delineate the role of m^6^A in innate immune response, we infected mouse RAW264.7 macrophage cells with different doses of gram-negative or -positive bacteria or human simplex virus type 1 (HSV-1), and examined the expression of innate immune response genes (Fig. S1). Infection with 10^7^ gram-positive bacteria *Corynebacterium diphtheriae*, 10^7^ *P. aeruginosa* or 1 multiplicity of infection (MOI) of HSV-1 induced maximum expression of innate immune response genes including colony stimulating factor 3 (CSF3), interleukin-1β (IL1β), transglutaminase 2 (TGM2), proto-oncogene tyrosine-protein kinase Src (SRC) under our experimental conditions (Fig. S1). Thus, we used these conditions in subsequent experiments.

We examined the protein levels of m^6^A “writers”, “erasers” and “readers” during bacterial and viral infections (Fig. 2A). The m^6^A “writer” protein METTL14 had marginal increases during *P. aeruginosa, C. diphtheriae* and wild-type (WT) HSV-1 infections while another m^6^A “writer” protein METTL3 had marginal increases during *P. aeruginosa* and *C. diphtheriae* infections (Fig. 2A). The eraser protein ALKBH5 also had slight increase during *C. diphtheriae* infection. Of the reader proteins examined, YTHDF1 had marginal increase during *C. diphtheriae* infection. However, YTHDF2 had significant increases during *C. diphtheriae* and *P. aeruginosa* infections by 3.89- and 4.21-fold, respectively, at 8 hpi (Fig. 2A). Since YTHDF2 mediates m^6^A-dependent RNA decay (23), we performed knockdown of YTHDF2 (Fig. 2B), and observed an upregulation of the DUSP1 transcript (Fig. 2C), which was also reflected in an increase of DUSP1 protein level (Fig. 2B). Knockdown of YTHDF2 almost doubled the half-life of DUSP1 transcript (Fig. 2D). Furthermore, YTHDF2 RNA immunoprecipitation reverse transcription quantitative real time PCR (RIP-qPCR) showed the binding of YTHDF2 protein to the DUSP1 RNA transcript, which was significantly increased at 4 and 6 hpi (Fig. 2E), correlating with the increased YTHDF2 protein level at these time points (Fig. 2A). Together, these results indicate that the upregulation of YTHDF2 protein promotes the degradation of DUSP1 transcript during *P. aeruginosa* infection.

**FIG2.**
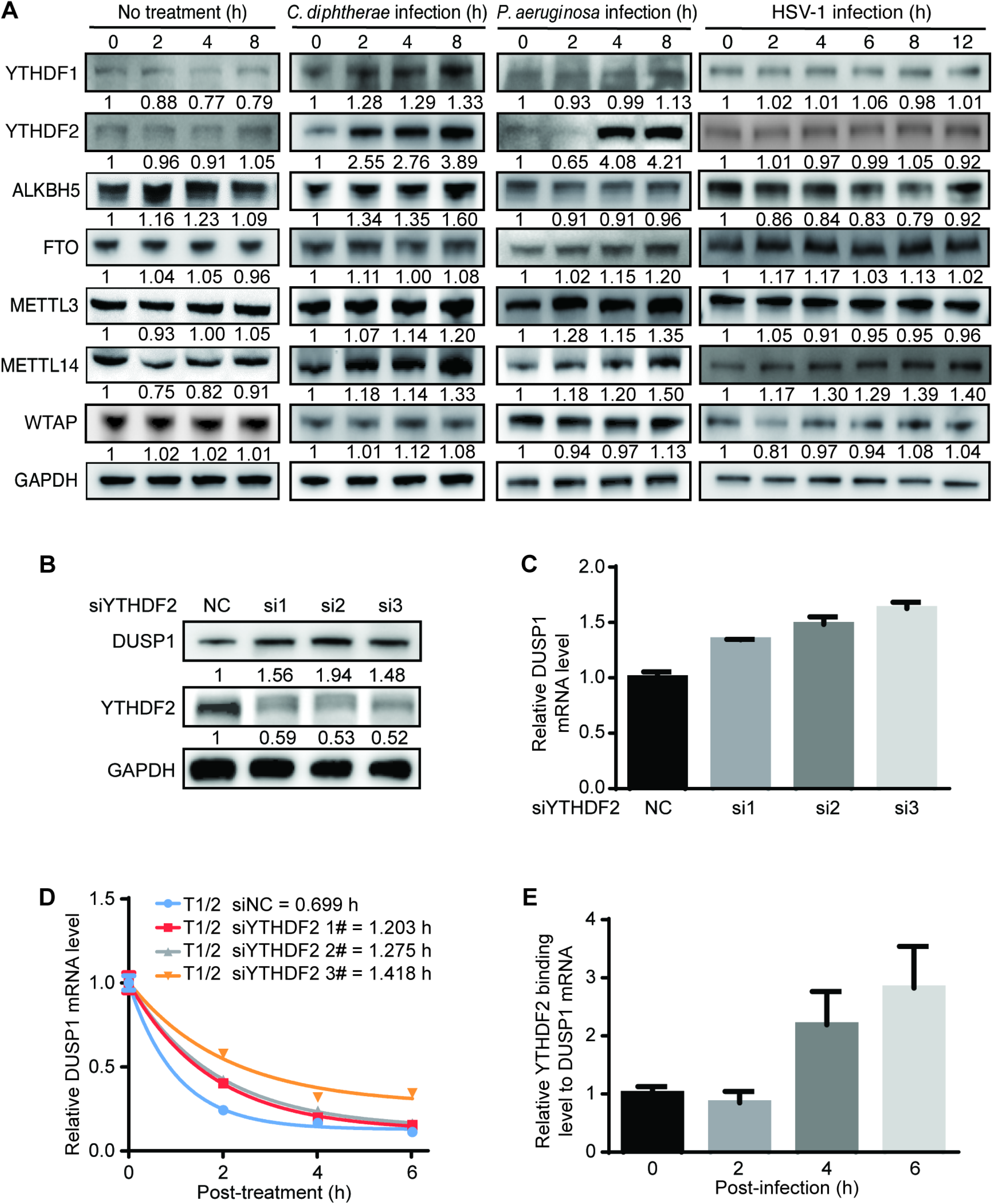
YTHDF2 mediates m^6^A-dependent DUSP1 transcript stability during bacterial and viral infections. (A) Protein levels of m^6^A writers METTL3, METTL14 and WTAP, erasers ALKBH5 and FTO, and readers YTHDF1 and YTHDF2 with or without infection by *C. diphtheriae*, *P. aeruginosa* or HSV-1 at the indicated time points examined by Western-blotting. (B) Examination of DUSP1 and YTHDF2 protein levels following YTHDF2 knockdown in RAW264.7 cells by Western-blotting. (C) Expression levels of DUSP1 transcript following YTHDF2 knockdown in RAW264.7 cells examined by RT-qPCR. (D) Alterations of half-lives of DUSP1 transcript following YTHDF2 knockdown in RAW264.7 cells during *P. aeruginosa* infection examined by RT-qPCR at the indicated time points following addition of 10 μg/ml actinomycin D. (E) Binding of YTHDF2 to DUSP1 transcript at the indicated time points following *P. aeruginosa* infection examined by RIP-qPCR.

### DUSP1 regulates p38 and JNK phosphorylation during bacterial and viral infections

As an important innate immune response gene, DUSP1 inactivates MAPKs by inhibiting their phosphorylation (11). Our results showed upregulation of DUSP1 transcript during bacterial and viral infections, which was reversed by m^6^A- and YTHDF2-mediated transcript degradation (Fig. 1 and 2). As expected, ERK, p38 and JNK MAPKs were activated at 2 hpi by *P. aeruginosa, C. diphtheriae*, and HSV-1 WT and ICP34.5 mutant viruses (Fig. 3A). We included the HSV-1 ICP34.5 mutant virus because the ICP34.5 protein has been shown to prevent the induction of innate immune genes during HSV-1 infection by directly inhibiting TBK1 activation and eIF2a function (24, 25). To determine whether DUSP1 regulated the activation of MAPKs during bacterial and viral infections, we performed DUSP1 knockdown.

**FIG 3.**
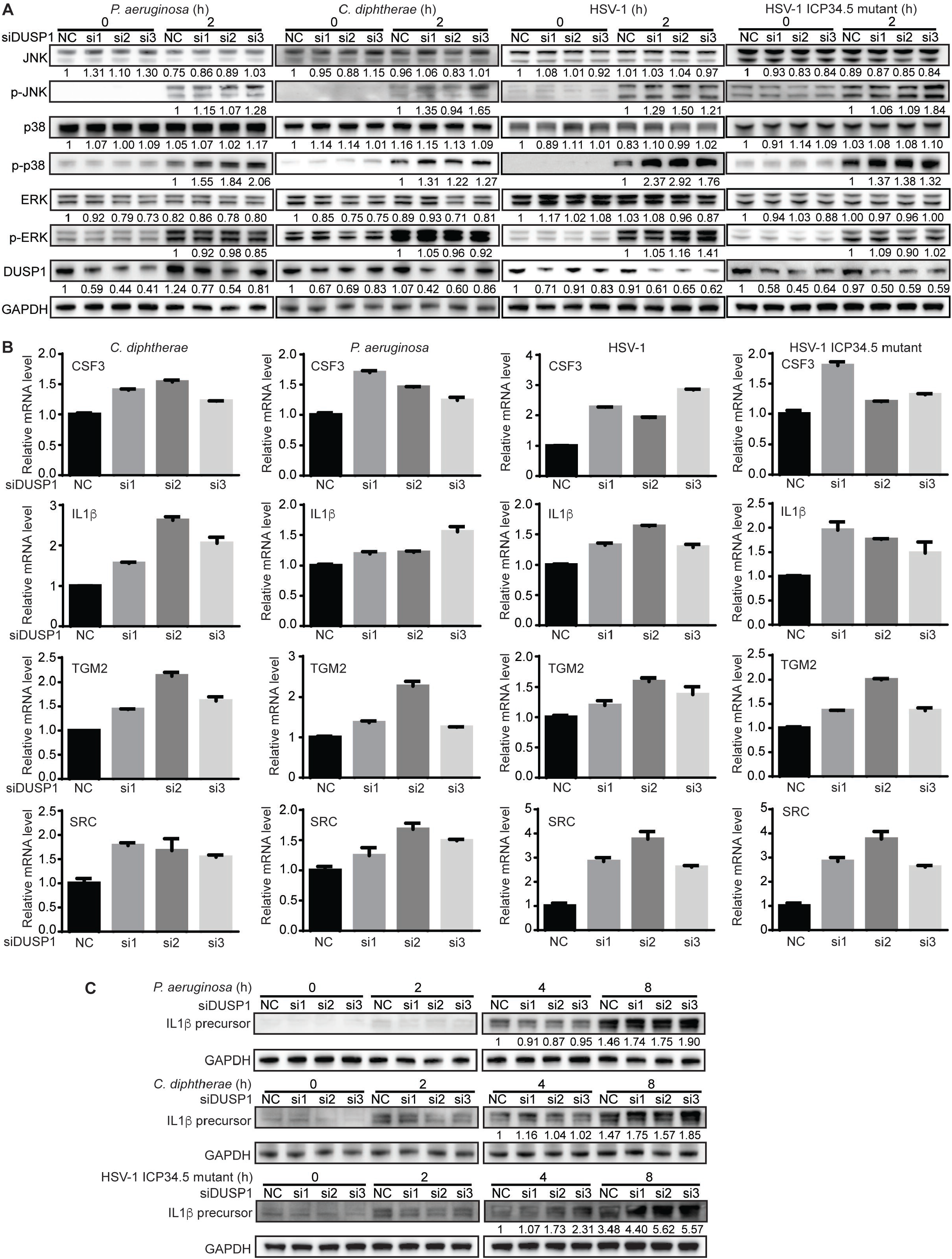
DUSP1 regulates p38 and JNK phosphorylation and expression of innate immune response genes during bacterial and viral infections. (A) DUSP1 knockdown enhanced the p38 and JNK phosphorylation during infection of *P. aeruginosa*, *C. diphtheriae*, or HSV-1 or HSV-1 ICP34.5 mutant virus. (B) DUSP1 knockdown enhanced the expression of IL1β, CSF3, TGM2 and SRC genes during infection of *P. aeruginosa, C. diphtheriae*, or HSV-1 or HSV-1 ICP34.5 mutant virus. (C) DUSP1 knockdown enhanced the protein level of IL1β during infection of *P. aeruginosa, C. diphtheriae* or HSV-1 ICP34.5 mutant virus.

Western-blotting results showed that the levels of phosphorylated p38 and JNK (p-p38 and p-JNK) were increased following DUSP1 knockdown during bacterial and viral infections (Fig. 3A). Activation of MAPKs can induce their downstream transcriptional factors including AP-1 and C/EBP, resulting in upregulation of target genes including numerous innate immune response genes (26). Consistent with the increased levels of p-p38 and p-JNK following DUSP1 knockdown, the levels of CSF3, IL1β, TGM2 and SRC transcripts were upregulated (Fig. 3B). We observed some variations of the effects of different DUSP1 siRNAs on the expression of IL1β, CSF3, TGM2 and SRC transcripts. These might be due to the different knockdown kinetics of these siRNAs. The IL1β protein level was also upregulated after DUSP1 knockdown during infections by *P. aeruginosa, C. diphtheriae*, and HSV-1 ICP34.5 mutant virus (Fig. 3C). However, upregulation of the IL1β protein was weak during WT HSV-1 infection and its increase was only marginal after DUSP1 knockdown (Fig. S2A), which was likely due to the inhibition of innate immune response by the HSV-1 ICP34.5 protein (24, 25). These results indicated that DUSP1 inhibited p-p38 and p-JNK activation to block innate immune response during bacterial and viral infections.

### ALKBH5 regulates p38 and JNK phosphorylation, and their downstream innate immune response genes during bacterial and viral infections

Since DUSP1 inactivated the p38 and JNK during bacterial and viral infections, and ALKBH5 knockdown reduced DUSP1 transcript stability by increasing m^6^A level, we examined ALKBH5 regulation of p38 and JNK activation. ALKBH5 knockdown increased the levels of p-p38 and p-JNK during infection by *P. aeruginosa, C. diphtheriae*, or HSV-1 WT or ICP34.5 mutant virus (Fig. 4A-4D). Some minor increase of p-ERK was also observed at 2 hpi of *C. diphtheriae*. Since the increased p-p38 and p-JNK levels could lead to enhanced activation of their downstream transcriptional factors, we examined *de novo* transcription of the target genes by performing nuclear run-on assay during *P. aeruginosa* infection. ALKBH5 knockdown indeed increased the transcriptional activities of IL1β, CSF3, TGM2 and SRC genes (Fig. 4E).

**FIG 4.**
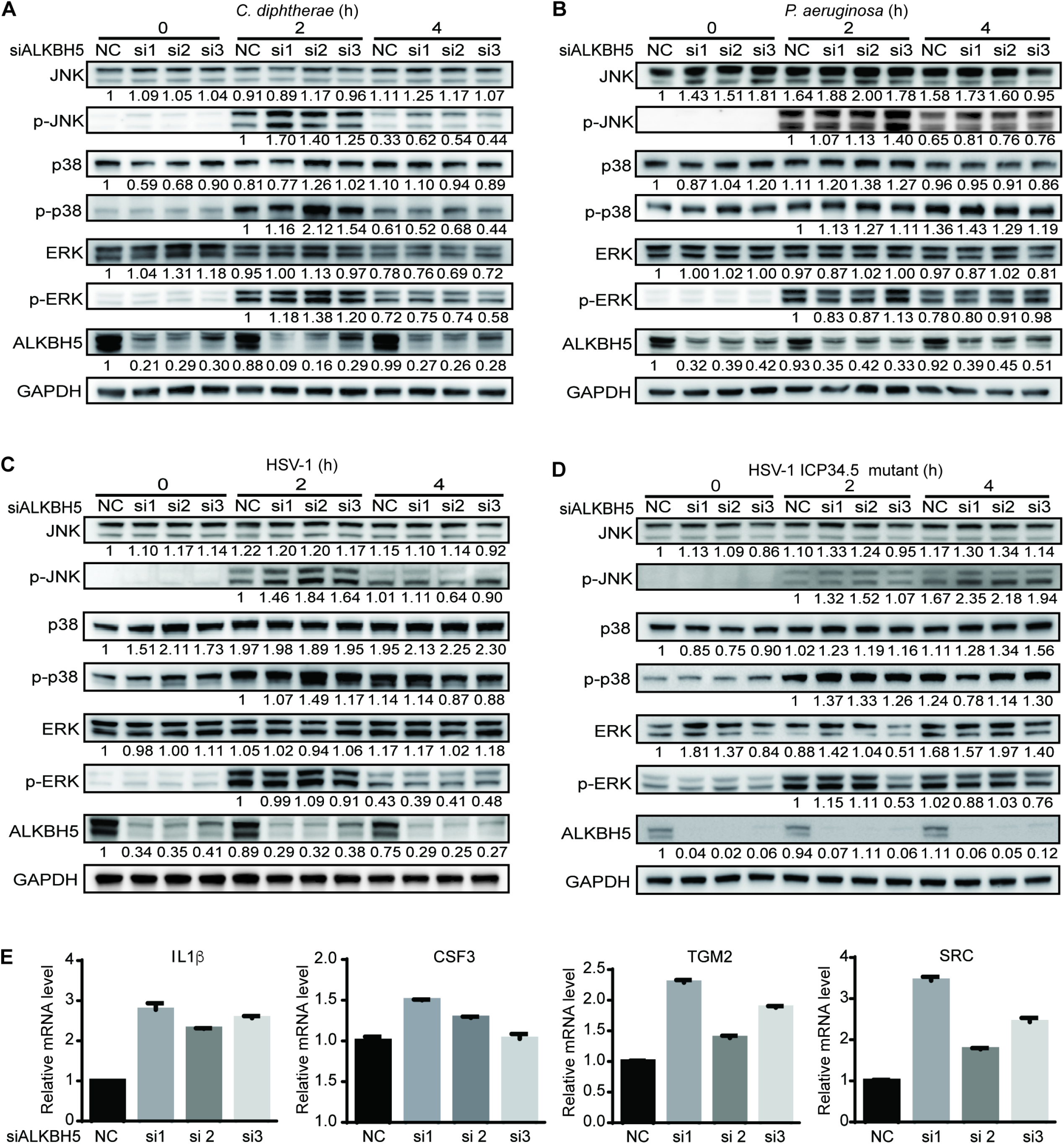
ALKBH5 regulates p38 and JNK phosphorylation and transcription of innate immune response genes during bacterial and viral infections. (A-D) ALKBH5 knockdown enhanced the p38 and JNK phosphorylation during infection of *C. diphtheriae* (A), *P. aeruginosa* (B), or HSV-1 (C) or HSV-1 ICP34.5 mutant virus (D). (E) *De novo* transcription of IL1β, CSF3, TGM2 and SRC genes following ALKBH5 knockdown at 2 hpi of *P. aeruginosa* examined by nuclear run-on assay. Cells treated with 4-thiouridine for 1 h after ALKBH5 knockdown were infected *P. aeruginosa* for 4 h and collected for nuclear run-on assay.

We further examined the role of ALKBH5 on the expression of innate immune response genes. ALKBH5 knockdown increased the levels of IL1β, CSF3, TGM2 and SRC transcripts during infection by *P. aeruginosa, C. diphtheriae*, or HSV-1 WT or ICP34.5 mutant virus (Fig. 5A). Similar to DUSP1 knockdown, we noticed variations of the effects of different ALKBH5 siRNAs on both the transcription and expression of IL1β, CSF3, TGM2 and SRC genes (Fig. 4E and 5). These might be due to the different knockdown kinetics of these siRNAs, which might impact the m^6^A level of DUSP1 transcript, DUSP1 expression level, and p-p38 and p-JNK levels, leading to variable transcription and expression levels of these downstream genes.

**FIG 5.**
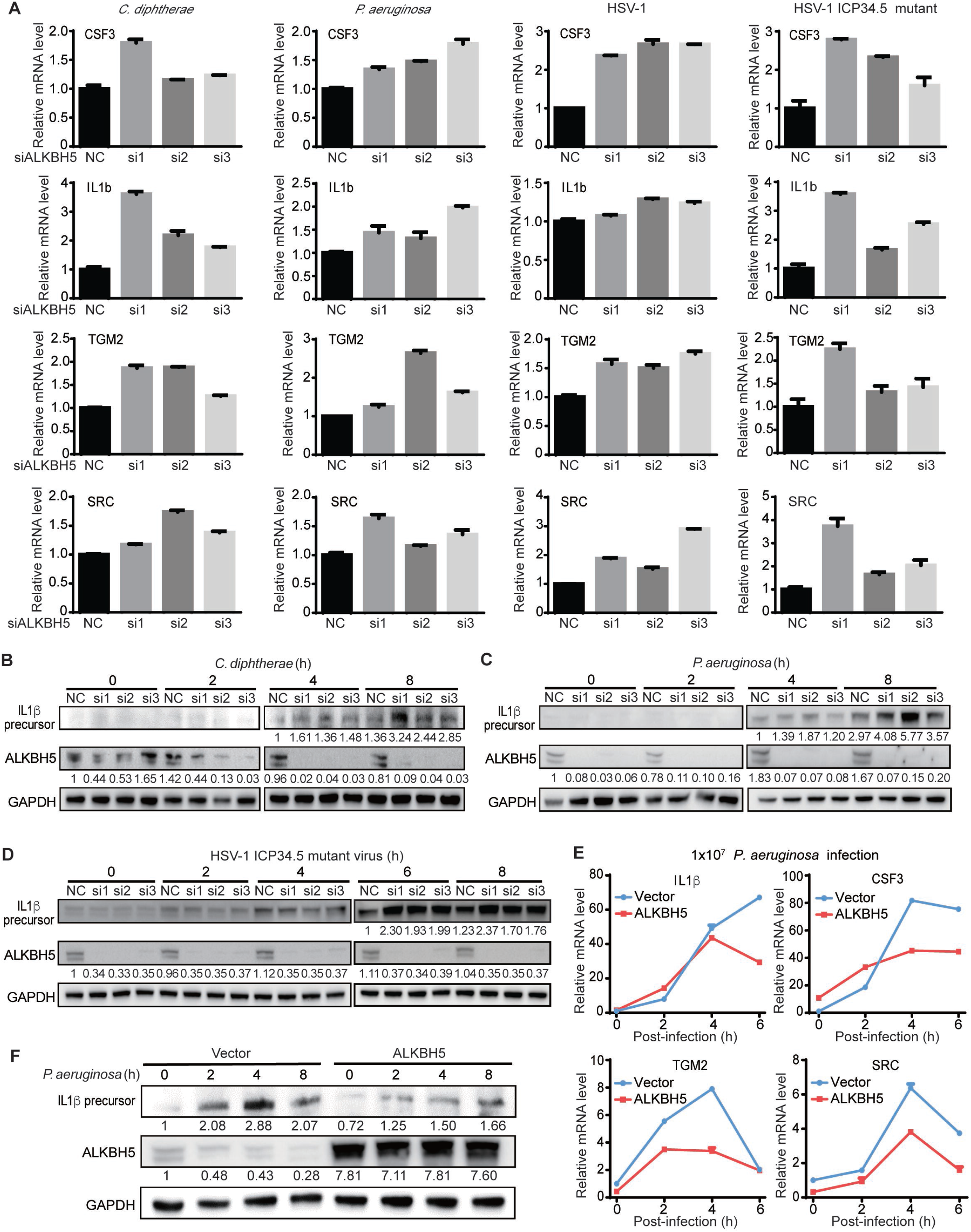
ALKBH5 regulates the expression of innate immune response genes during bacterial and viral infections. (A) ALKBH5 knockdown enhanced the expression levels of IL1β, CSF3, TGM2 and SRC transcripts during *P. aeruginosa, C. diphtheriae*, or HSV-1 or HSV-1 ICP34.5 mutant virus infection. (B–D) ALKBH5 knockdown enhanced the protein level of IL1β during infection by *C. diphtheriae* (B), *P. aeruginosa* (C), or HSV-1 ICP34.5 mutant virus (D). (E) ALKBH5 overexpression inhibited the expression of IL1β, CSF3, TGM2 and SRC genes during *P. aeruginosa* infection measured by RT-qPCR. (F) ALKBH5 overexpression inhibited the protein level of IL1β during *P. aeruginosa* infection measured by Western-blotting.

The protein level of IL1β was also upregulated after ALKBH5 knockdown during infections by *P. aeruginosa, C. diphtheriae*, and HSV-1 ICP34.5 mutant virus (Fig. 5B-5D). However, the upregulation of IL1β protein was marginal during WT HSV-1 infection (Fig. S2B). In contrast, overexpression of ALKBH5 reduced the levels of IL1β, CSF3, TGM2 and SRC transcripts (Fig. 5E), and downregulated the IL1β protein level (Fig. 5F) during *P. aeruginosa* infection. It was interesting that the reduced expression of the four transcripts had different kinetics following overexpression of ALKBH5 (Fig. 5E). The effect of ALKBH5 overexpression was observed for TGM2 and SRC transcripts at as early as 2 hpi, which disappeared by 6 hpi. However, the effect was not observed for CSF3 until 4 hpi and for IL1β until 6 hpi. It is possible that the promoters of these genes might endow them with different kinetics in response to the activation of p38 and JNK pathways.

Because our results showed an important role of ALKBH5 in regulating innate immune response, we further examined the impact of ALKBH5 knockdown on HSV-1 replication. ALKBH5 knockdown reduced the replication of HSV-1 WT or ICP34.5 mutant virus (Fig. S3). These results are in agreement with those of a previous study showing reduced HSV-1 replication after ALKBH5 knockout (14).

In conclusion, bacterial and viral infections activate MAPKs to induce innate immune response genes as well as a negative regulator of MAPKs, DUSP1, to avoid excessive innate immune response. At the same time, numerous m^6^A writer proteins and reader protein YTHDF2 are induced, leading to hyper-methylation of DUSP1 transcript, which is targeted for YTHDF2-mediated degradation. This mechanism of fine-tuned activation of MAPKs optimizes the induction of innate immune response genes during pathogenic infections (Fig. 6).

**FIG 6.**
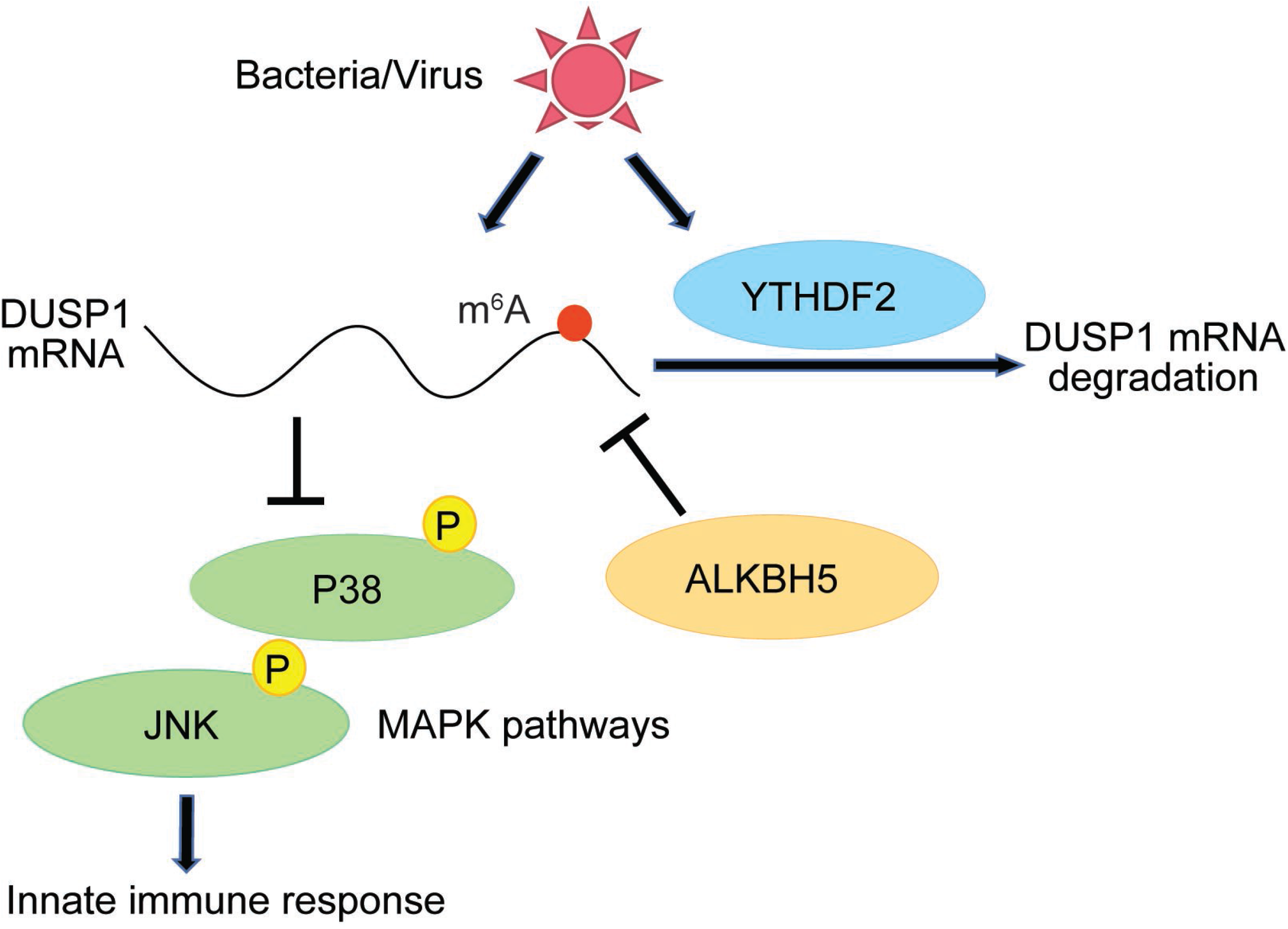
A working model of regulation of DUSP1, MAPKs and innate immune response genes by m^6^A and m^6^A-related proteins YTHDF2 and ALKBH5 during pathogenic infections.

## Discussion

The innate immune system is a complex cellular and molecular network in mammalian cells that serves as the first line of defense against pathogenic infections and is regulated by diverse cellular pathways (1). DUSP1 is a critical regulator of MAPK pathways serving as a negative feedback mechanism to prevent excessive activation of these pathways (27, 28). In the context of pathogenic infections, activation of MAPKs induces the expression of innate immune response genes as well as DUSP1, which prevents overreactive immune response (29–31). Our results showed that DUSP1 transcript was indeed induced during bacterial and viral infections together with the activation of the ERK, JNK and p38 MAPK pathways. At the same time, the m^6^A level of DUSP1 transcript was significantly increased. During these processes, we only observed marginal increases of m^6^A writer proteins METTL3 and METTL14 and no decrease of m^6^A eraser proteins ALKBH5 and FTO, suggesting that the observed m^6^A increase in the DUSP1 transcript likely depended on preexisting writer proteins. Interestingly, despite the increased expression of DUSP1 transcript during bacterial and viral infections, we failed to detect an increase of DUSP1 protein. It is unclear whether the increased DUSP1 transcript m^6^A might affect its translation. In addition, it is unclear why the m^6^A level is only increased in some *de novo* transcribed transcripts but not others. The specific mechanism involved in this selection process might deserve further investigations. Nevertheless, the results of ALKBH5 knockdown experiments revealed that the m^6^A increase in the DUSP1 transcript targeted it for YTHDF2-mediated degradation. Importantly, YTHDF2 was significantly induced during bacterial infections, which maximized its negative regulation of DUSP1 transcript stability. Taken together, these results suggest that m^6^A and YTHDF2 are involved in fine-tuning the expression of DUSP1 protein, an important regulator of innate immunity, during pathogenic infections.

The observed induction of YTHDF2 protein is consistent with results from another study showing LPS induction of YTHDF2 expression (32). Interestingly, there was no obvious change of YTHDF2 protein following HSV-1 infection, indicating possible involvement of the bacteria-associated pattern recognition receptors in the induction of YTHDF2 protein. However, it is possible that HSV-1 infection might have a YTHDF2 induction kinetic that is different from those of bacterial infections. Alternatively, HSV-1 might have evolved to prevent YTHDF2 induction as a mechanism to counter innate immune response.

Our results showed that m^6^A- and YTHDF2-mediated degradation of DUSP1 transcript resulted in enhanced activation of p38 and JNK. Both p38 and JNK pathways activate transcriptional factors such AP-1 and C/EBP that are essential for the expression of innate immune response genes. Indeed, knockdown DUSP1 or m^6^A eraser ALKBH5 enhanced the expression of innate immune response genes including IL1β, CSF3, TGM2 and SRC during bacterial or viral infections. We observed robust induction of the IL1β precursor by HSV-1 ICP34.5 mutant but not WT virus (Fig. S2). It has been reported that the ICP34.5 protein can directly inhibit TBK1 and eIF2a proteins to prevent the induction of innate immune genes during HSV-1 infection (24, 25). Interestingly, activated MAPK pathways can promote HSV-1 viral replication by activating downstream transcriptional factors (33, 34). However, we showed that ALKBH5 knockdown inhibited HSV-1 replication, which was likely due to m^6^A-mediated downregulation of DUSP1 and subsequent activation of MAPK pathways resulting in the induction of innate immune response. However, it is also possible that ALKBH5 and m^6^A might regulate HSV-1 replication through another mechanism in addition to targeting DUSP1 transcript for degradation and activating MAPK pathways.

We have previously shown that a set of innate immune response genes are subjected to m^6^A modification and might be directly regulated by m^6^A while another set of innate immune response genes might be indirectly regulated by m^6^A during bacterial and viral infections (13). In the current work, we have provided an example of m^6^A and YTHDF2 indirect regulation of innate immune response genes by mediating the stability of DUSP1 transcript. In fact, DUSP1 is under the tight control of m^6^A and YTHDF2 during bacterial and viral infections. It can be speculated that other DUSP genes, which are involved in diverse cellular functions, could also be regulated by m^6^A and m^6^A-related proteins, and therefore deserve further investigations.

## Material and methods

### Bacteria, viruses, and cells

*P. aeruginosa* and *C. diphtheriae* were purchased from ATCC. Herpes simplex virus type 1 (HSV-1) F strain and HSV-1 ICP34.5 mutant virus were obtained from Dr. Bernard Roizman (The University of Chicago, Chicago, IL). The ICP34.5 mutant virus (R3616) was generated by deleting the 1 kb fragment containing both copies of the gamma 34.5 gene between the BstEII and Stu I sites from the F strain HSV-1 genome (35). RAW 264.7 cells were purchased from ATCC and cultured following the instructions of the vendor.

### Bacteria and virus infection

RAW 264.7 cells at 4×10^5^ cells per mL were infected with *P. aeruginosa* or *C. diphtheriae* at 10^7^ per mL, or with HSV-1 WT or ICP34.5 mutant virus at 1 MOI. Cells were harvested at the indicated time points.

### m^6^A-immunoprecipitation (m^6^A-IP)

Isolation of m^6^A-containing fragments was performed as previously described (13, 36). Briefly, total RNA was extracted from cells using TRI Reagent (T9424-200ML, Sigma-Aldrich) and fragmented using RNA fragmentation kit (AM8740, ThermoFisher). Successful fragmentation of RNA with sizes close to 100 nucleotides was validated using bioanalyzer (2100 Bioanalyzer Instrument, Agilent). Anti-m^6^A antibody (10 μg) (202-003, Synaptic Systems) was incubated with 30 μl slurry of Pierce Protein A Agarose beads (20365, ThermoFisher) by rotating in 250 μl PBS at 4 °C for 3 h. The beads were washed three times in cold PBS followed by one wash in IP buffer containing 10 mM Tris-HCl at pH 7.4, 150 mM NaCl, and 1% Igepal CA-630 (I8896-50ML, Sigma-Aldrich). To isolate the m^6^A-containing fragments, 120 μg of fragmented total RNA was added to the antibody-bound beads in 250 μl IP buffer supplemented with RNasin Plus RNase inhibitor (PRN2615, Promega), and the mixture was incubated at 4 °C for 2 h. The beads were washed seven times with 1 ml IP buffer and eluted with 100 μl IP buffer supplemented with 6.67 mM of m^6^A salt (M2780, Sigma-Aldrich) at 4 °C for 1 h. A second elution was carried out and the eluates were pooled together before purification by ethanol 70% precipitation.

### siRNA knockdown

siRNA silencing was performed by transfecting 2.5 pmol of each siRNA per well in a 12-well plate into the RAW264.7 cells using Lipofectamine RNAi Max (13778150, ThermoFisher) according to manufacturer’s instructions. Two days after transfection, the cells were monitored for knockdown efficiency of the target gene by RT-qPCR and Western-blotting. siRNAs purchased from Sigma-Aldrich are as follows: DUSP1 si1: SASI_Mm02_00322441; DUSP1 si2: SASI_Mm01_00056586; DUSP1 si3: SASI_Mm01_00056587; ALKBH5 si1: SASI_Mm01_00106232; ALKBH5 si2: SASI_Mm02_00344968; ALKBH5 si3: SASI_Mm01_00106233; and siNegative Control (NC): Sigma siRNA Universal Negative Control #1 (SIC001-10NMOL).

### RNA stability assay

Actinomycin D (10 μg/ml) (A9415-2MG, Sigma-Aldrich) was added to cells to inhibit transcription. RNA was isolated at 0, 2, 4 and 6 h after actinomycin D treatment using Trizol and the transcripts were quantified by RT-qPCR.

### RT-qPCR for gene expression, RIP-qPCR for YTHDF2 RNA binding quantification and MeRIP-qPCR for m^6^A-seq validation

Total RNA was isolated with TRI Reagent (T9424-200ML, Sigma-Aldrich) according to the manufacturer’s instructions. Reverse transcription was performed with 1 μg of total RNA using Maxima H Minus First Strand cDNA Synthesis Kit (Cat.# K1652, ThermoFisher). Quantitative PCR was done using SsoAdvanced Universal SYBR Green Supermix (1725271, Bio-Rad). Relative gene expression levels were obtained by normalizing the cycle threshold (CT) values to yield 2^-ΔΔCt^ values. For validation of m^6^A-seq, eluted or input mRNA was subjected to RT-qPCR. Fold enrichment was obtained by calculating the 2^-ΔCt^ value of eluate in relative to the input sample. The primers used for gene expression are:

5’CTGGTGGGTGTGTCAAGCAT3’ (forward) and

5’GAGGCAGTTTCTTCGCTTGC3’ (reverse) for DUSP1; and

5’CCCTGAAGTACCCCATTGAA3’ (forward) and

5’GGGGTGTTGAAGGTCTCAAA3’ (reverse) for β-actin; and

5’GAGTGTGGATCCCAAGCAAT3’ (forward) and

5’ACGGATTCCATGGTGAAGTC3’ (reverse) for IL1β; and

5’CCGGTACCCTCTCCTGTTGTGTTTA3’ (forward) and

5’AACTCGAGCTAAAAAGGAGGACGGC3’ (reverse) for CSF3; and

5’AAGAGCTCCAAACAAGGTCTGCCTT3’ (forward) and

5’AACTCGAGACGTGCCATATAAGCAC3’ (reverse) for TGM2; and

5’AAGGTACCCTGCCAGGCCAGACCAA3’ (forward) and

5’AACTCGAGCCAGCCTTGACCCTGAG3’ (reverse) for SRC; and

5’ACGGTTTACTACGCCGTGTT3’ (forward) and

5’TGTAGGGTTGTTTCCGGACG3’ (reverse) for US6; and

5’GACGAACATGAAGGGCTGGA’ (forward) and

5’CGACCTGTTTGACTGCCTCT3’ (reverse) for VP16; and

5’CCCACTATCAGGTACACCAGCTT3’ (forward) and

5’CTGCGCTGCGACACCTT3’ (reverse) for ICP0; and

5’GCATCCTTCGTGTTTGTCATTCTG3’ (forward) and

5’GCATCTTCTCTCCGACCCCG3’ (reverse) for ICP27.

### Western-blotting

Protein samples were lysed in Laemmli buffer, separated by SDS-PAGE and transferred to a nitrocellulose membrane (37). The membrane was blocked with 5% milk and then incubated with primary antibody to GAPDH (5174s, CST), p38 (8690S, CST), p-p38 (4511S, CST), ERK (4695S, CST), p-ERK (4370S, CST), JNK (9252S, CST), p-JNK (4668S, CST), DUSP1 (NBP2-67909, Novus), IL1β (AB-401-NA, R&D system), or ALKBH5 (HPA007196, Sigma) overnight at 4 °C. The membrane was washed with TBS-Tween (TBS-T) and probed with a secondary antibody conjugated to horseradish peroxidase (HRP). After further washing with TBS-T, the blot was visualized using SuperSignal^™^ West Femto Maximum Sensitivity Substrate (34096, Thermo) and imaged on a ChemiDoc^™^ MP Imaging System (12003154, Bio-Rad).

### Nuclear run-on assay

Nuclear run-on assay was conducted as previously described (38).

### RNA immunoprecipitation (RIP) assay

RIP assay was conducted as previously described (39).

## SUPPLEMENTAL MATERIAL

Supplemental material is available online only.

FIG S1, PDF file, 0.5 MB.

FIG S2, PDF file, 1 MB.

FIG S3, PDF file, 0.8 MB.

## ACKNOWLEDGMENTS

This work was supported by grants from the National Institutes of Health (CA096512 and CA124332 to S.-J. Gao), and in part by award P30CA047904.

We thank members of Dr. Shou-Jiang Gao’s laboratory for technical assistance and helpful discussions.

J.F. performed most of the experiments. W.M., L.P.C., X.Q.Z. and A.M. performed a subset of experiments. Y.F.H. performed the bioinformatic analysis. W.Y. provides the HSV-1 wild-type and mutant viruses. J.F. and S.J.G. prepared the manuscript. S.J.G. planned, managed, and supervised the study, and secured funding.

## SUPPLEMENTAL MATERIAL

**FIG S1** Expression of CSF3, IL1β, TGM2 and SRC transcripts following infection with different doses of *C. diphtheriae, P. aeruginosa* or HSV-1 at the indicated time points examined by RT-qPCR.

**FIG S2** Protein level of IL1β precursor following knockdown of DUSP1 (A) or ALKBH5 (B) at different time points following HSV-1 infection.

**FIG S3** ALKBH5 knockdown inhibits HSV-1 replication. RAW264.7 cells transfected with ALKBH5 shRNAs or a scrambled control (NC) were infected with 1 MOI of HSV-1 for 48 h, and the supernatants were collected for plaque assay to determine the viral titers.

